## Supplemental Figure 1-1 for "*In vivo* functional diversity of midbrain dopamine neurons within identified axonal projections"

A

FG-labeling detection (intrinsic FG signal)

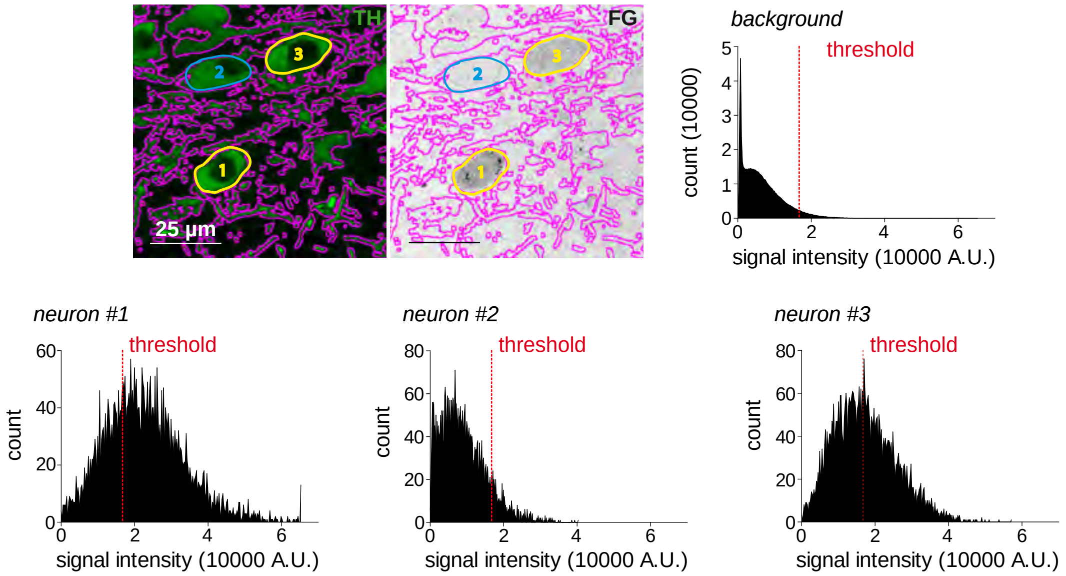


| TH neuron # | mean (TH+ neuron) | mean (TH-free BG) | BG SD | threshold | FG-labelled? |
|---|---|---|---|---|---|
| 1 | 22899 | 6704 | 5013 | 16730 | ✔ |
| 2 | 9119 | 6704 | 5013 | 16730 | ✘ |
| 3 | 17385 | 6704 | 5013 | 16730 | ✔ |

B

FG-labeling detection (FG-Antibody)

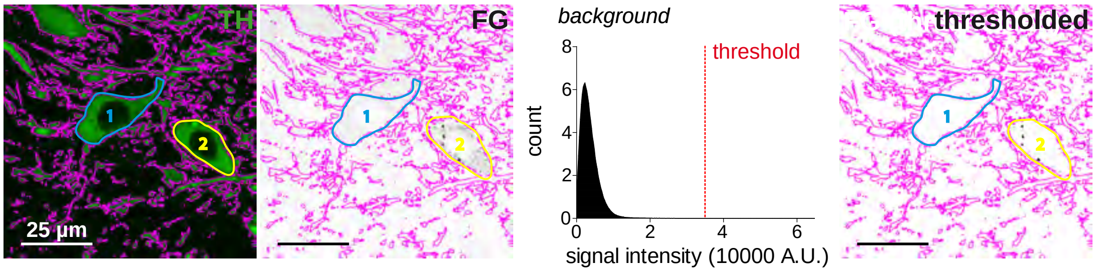


| TH neuron # | mean (TH-free BG) | BG SD | threshold | positive marks | FG-labelled? |
|---|---|---|---|---|---|
| 1 | 3720 | 3037 | 34973 | 0 | ✘ |
| 2 | 3720 | 3037 | 34973 | >5 | ✔ |
