## Supplementary figures and images for "*In vivo* functional diversity of midbrain dopamine neurons within identified axonal projections"

### Supplemental Figure 3-1

##### Supplemental Figure 3-1

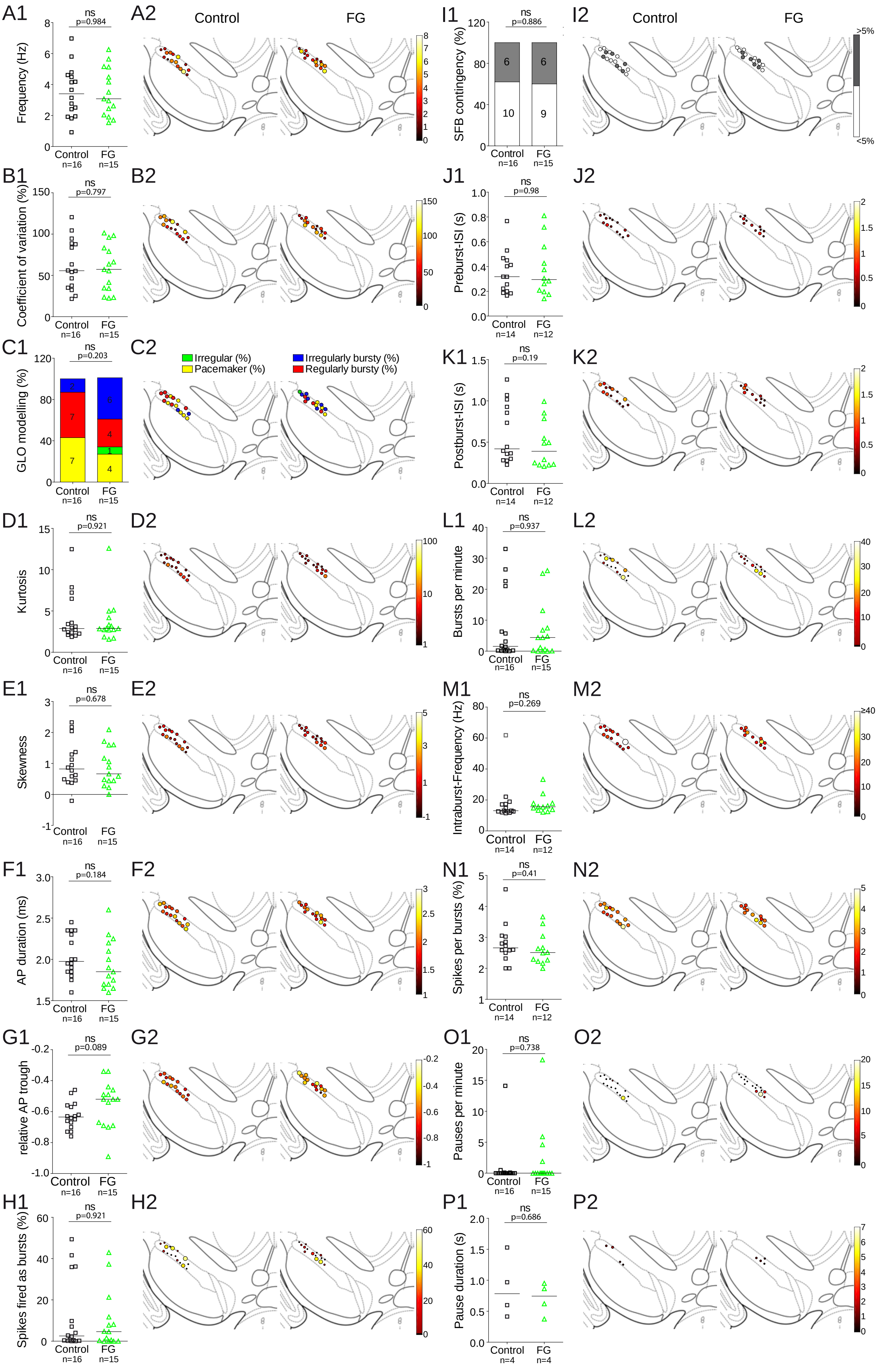

### Supplemental Figure 4-1

#### Supplemental Figure 4-1

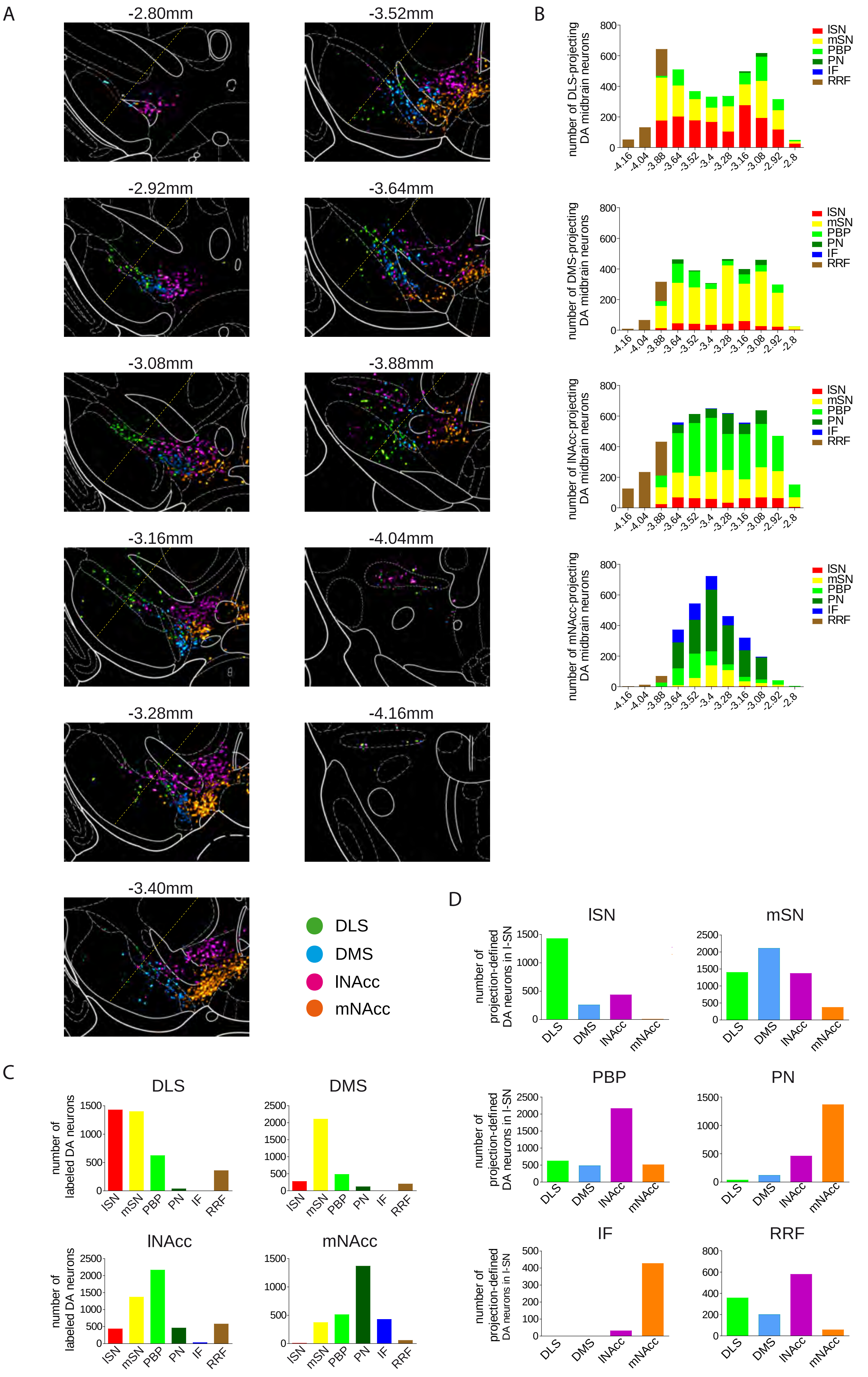

### Supplemental Figure 4-2

Supplemental Figure 4-2

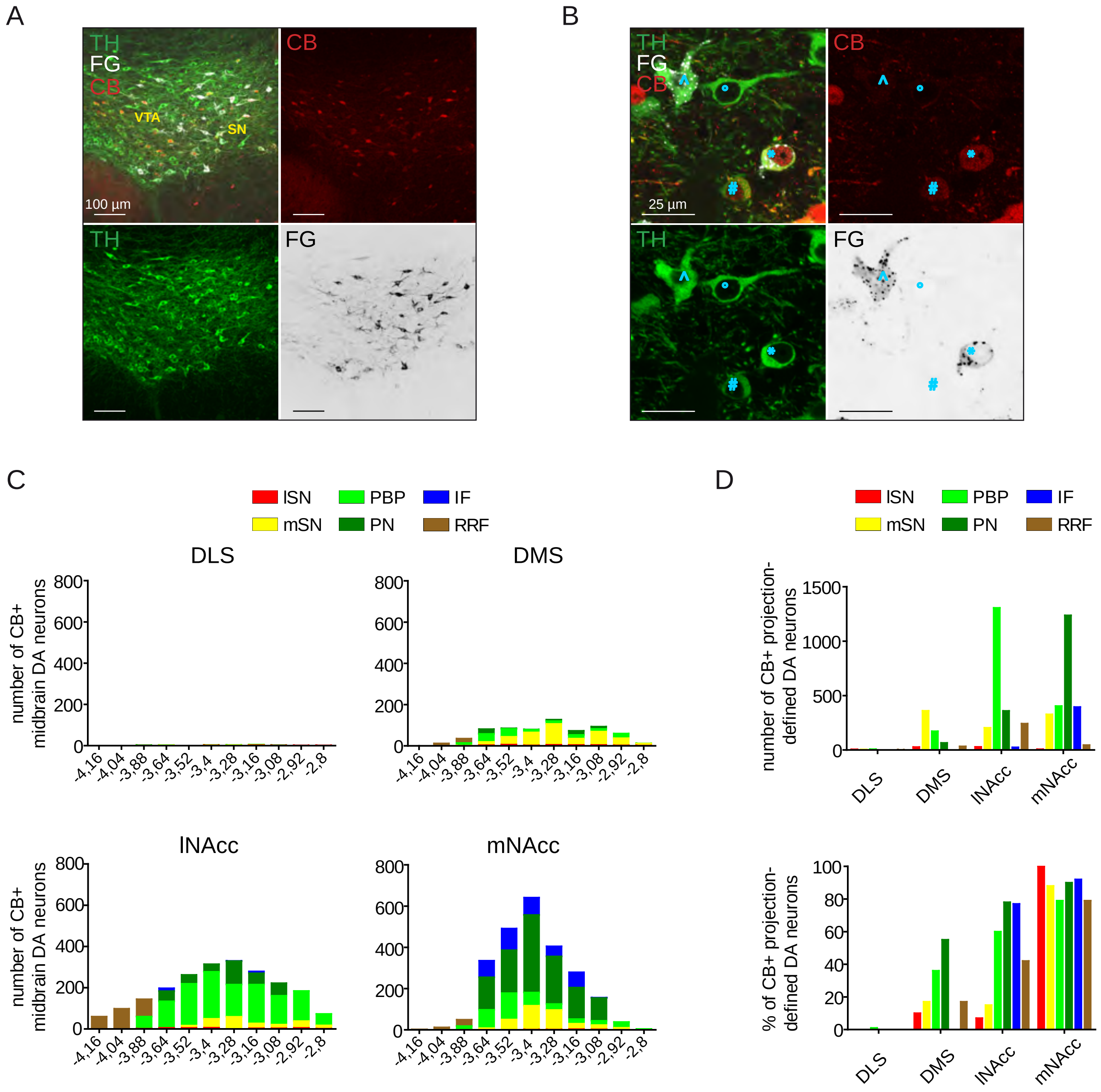

### Supplemental Figure 5-1

#### Supplemental Figure 5-1

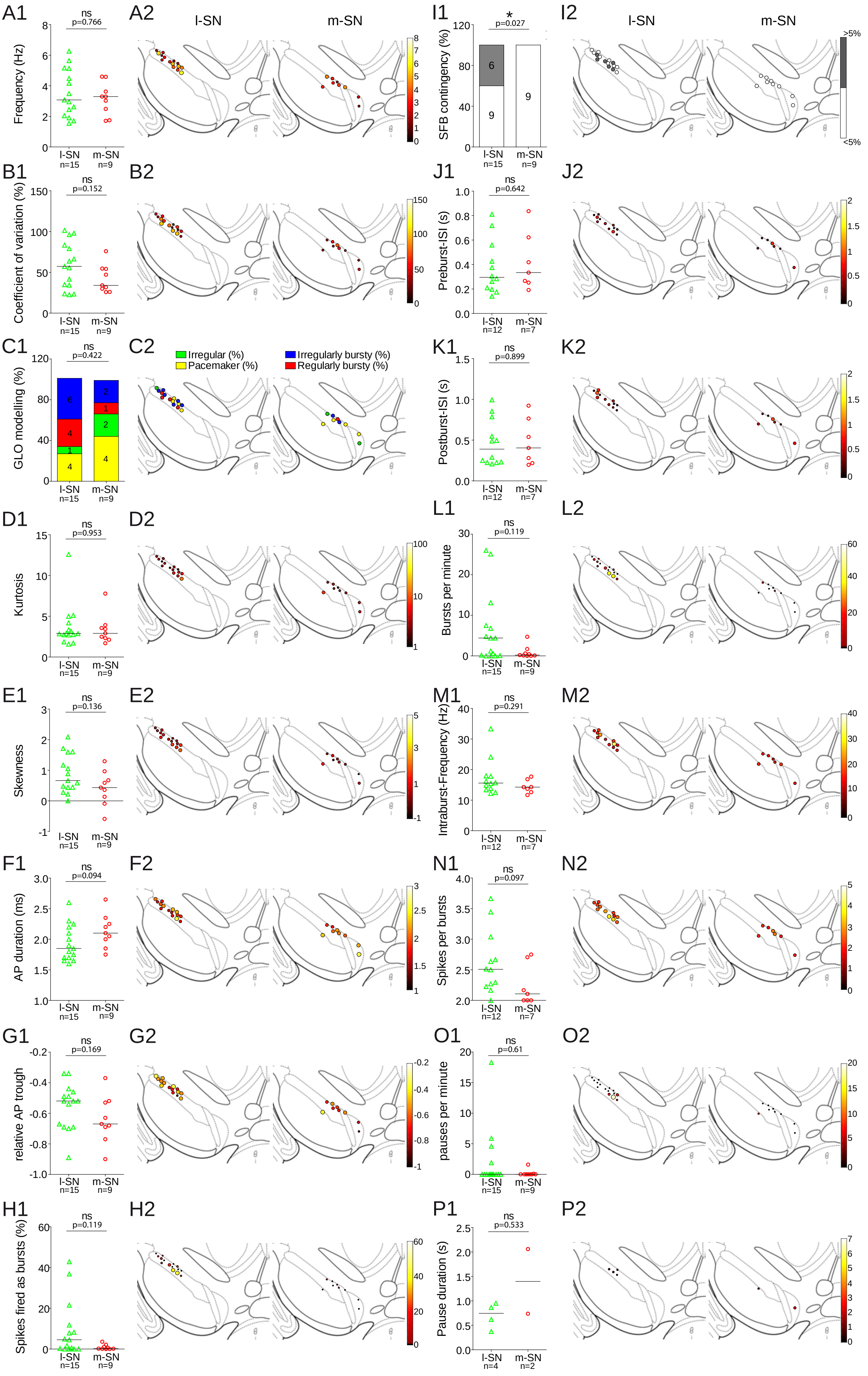

### Supplemental Figure 6-1

Supplemental Figure 6-1

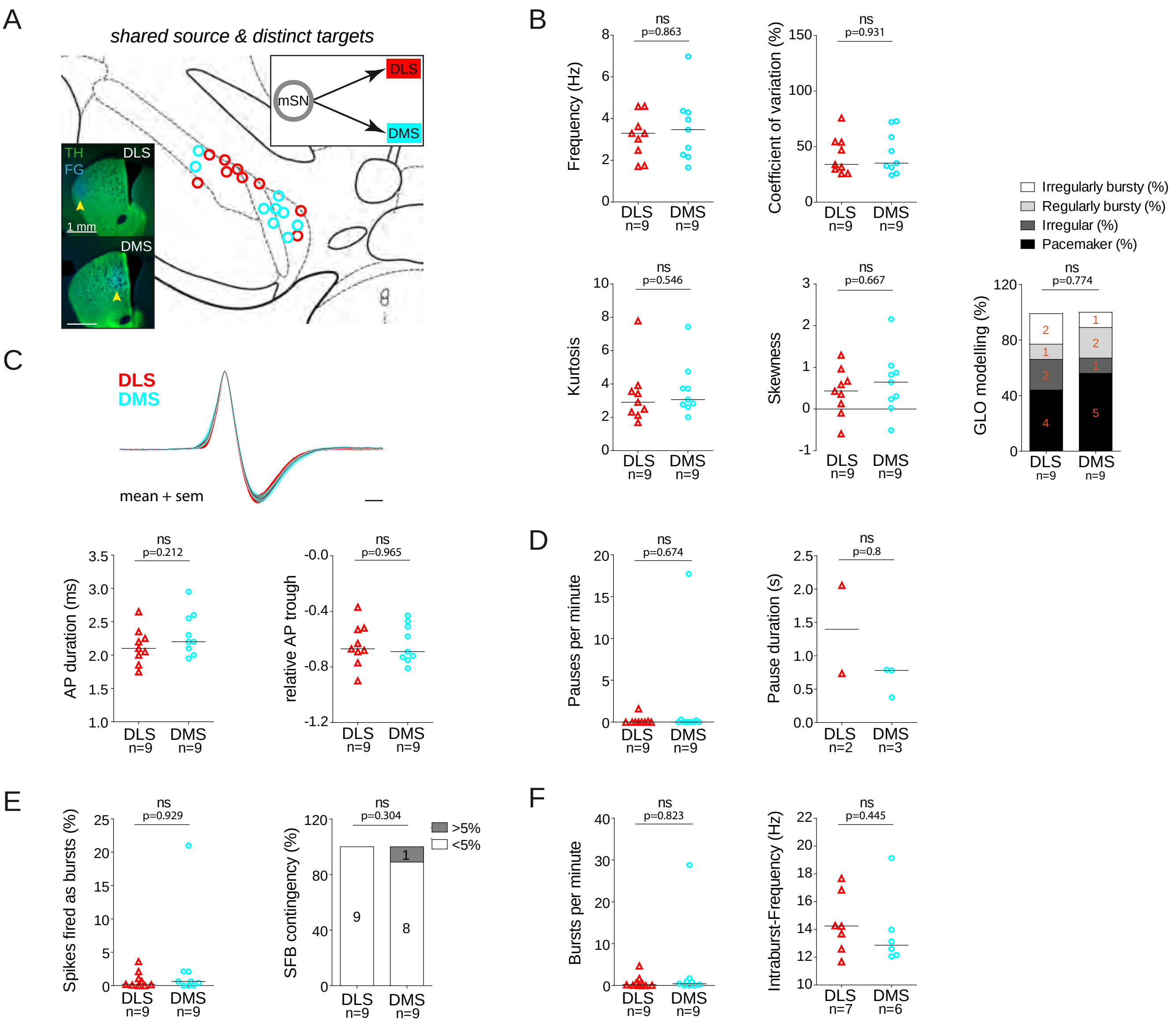

### Supplemental Figure 6-2

Supplemental Figure 6-2

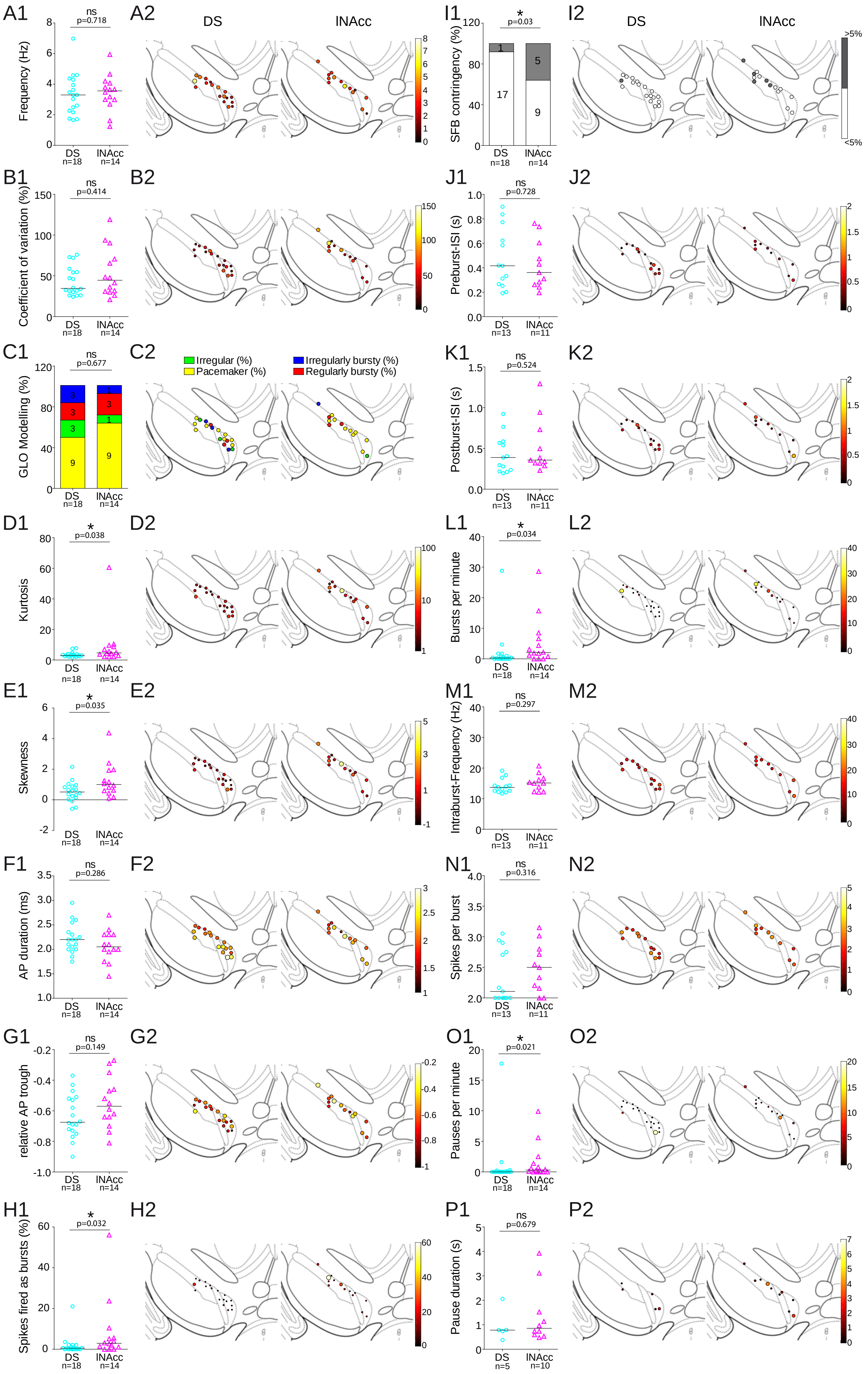

### Supplemental Figure 6-3

#### Supplemental Figure 6-3

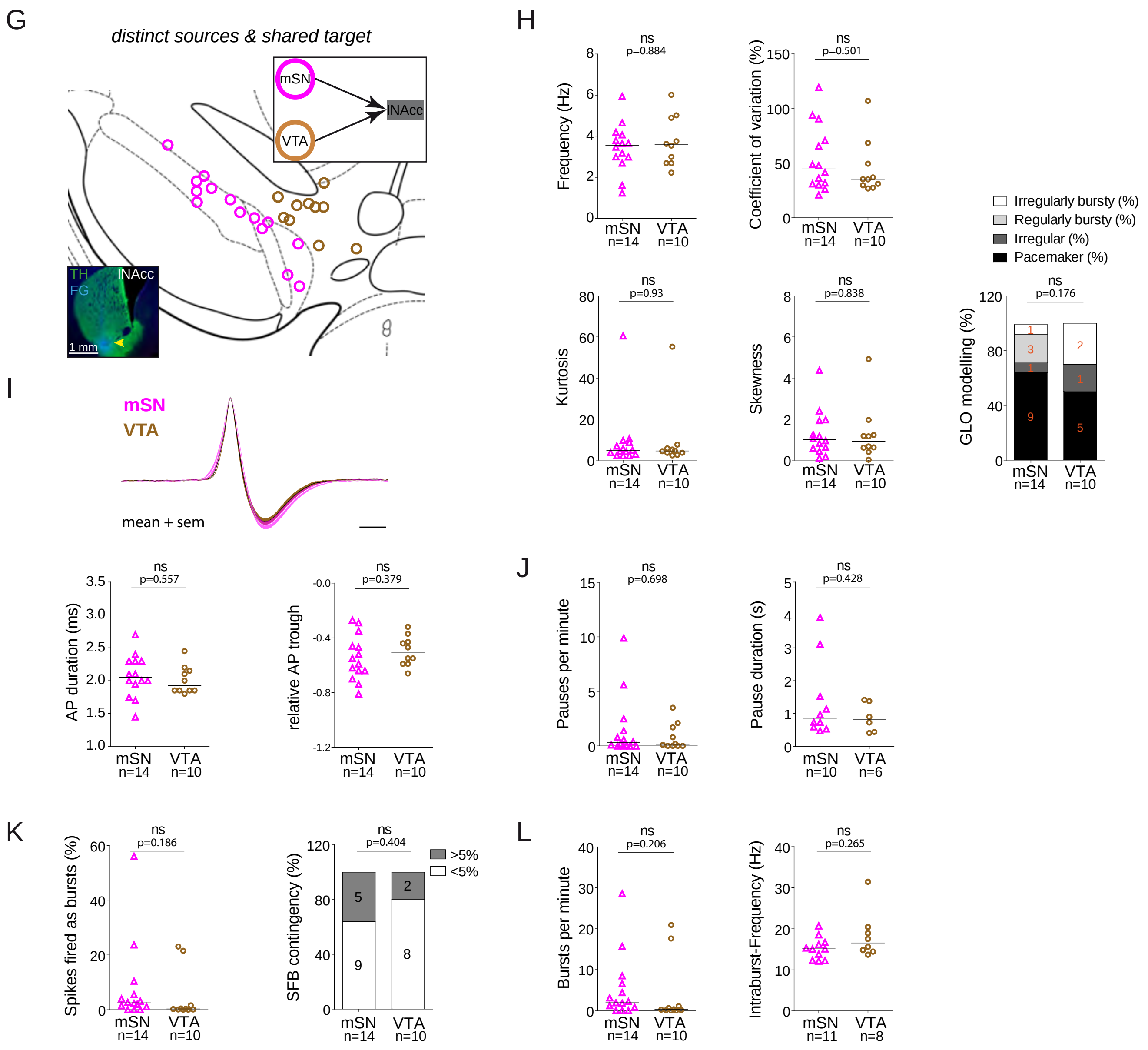

### Supplemental Figure 7-1

Supplemental Figure 7-1

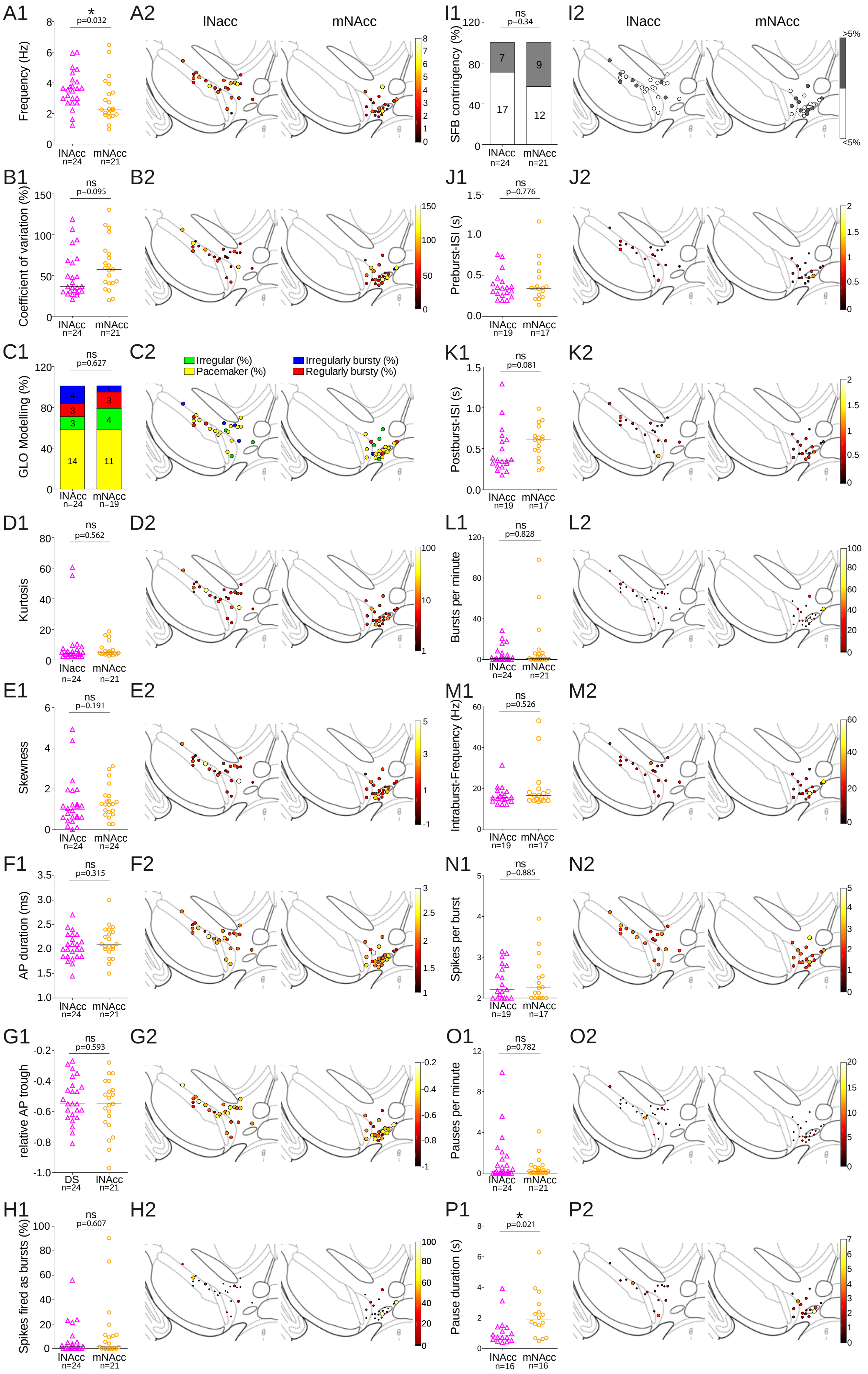

### Supplemental Figure 7-2

Supplemental Figure 7-2

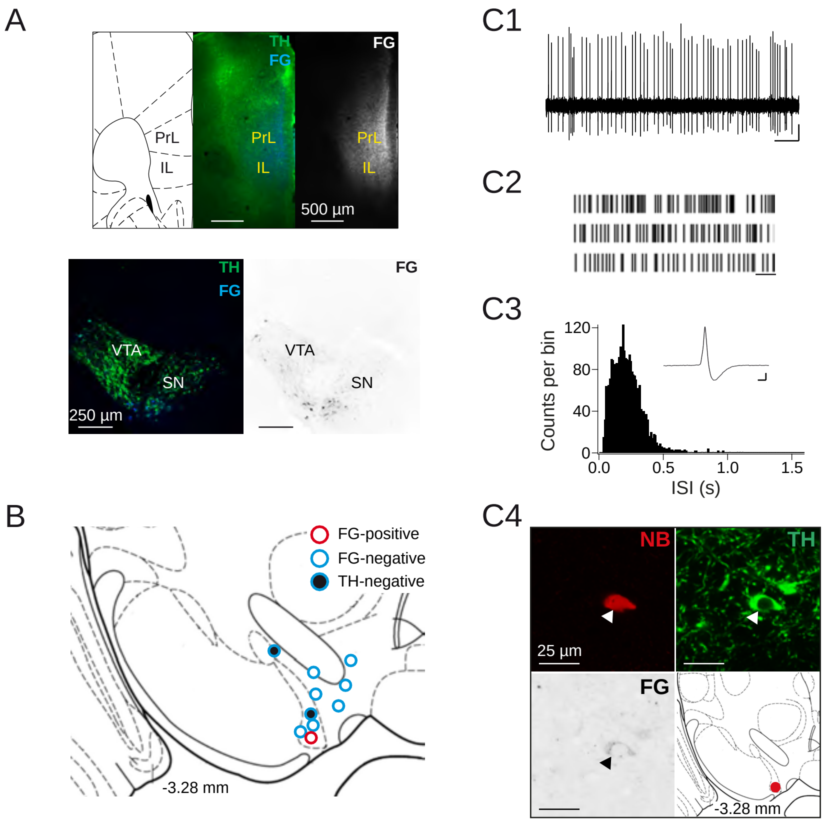
